# Identification of QTLs for dynamic and steady state photosynthetic traits in a barley mapping population

**DOI:** 10.1101/2020.08.09.243352

**Authors:** William T. Salter, Si Li, Peter M. Dracatos, Margaret M. Barbour

## Abstract

Enhancing the photosynthetic induction response to fluctuating light has been suggested as a key target for improvement in crop breeding programs, with the potential to substantially increase whole canopy carbon assimilation and contribute to crop yield potential. Rubisco activation may be the main physiological process that will allow us to achieve such a goal. In this study, we phenotypically assessed the rubisco activation rate in a doubled haploid (DH) barley mapping population [131 lines from a Yerong/Franklin (Y/F) cross] after a switch from moderate to saturating light. Rates of rubisco activation were found to be highly variable across the mapping population, with a median activation rate of 0.1 min^−1^ in the slowest genotype and 0.74 min^−1^ in the fastest genotype. A QTL for rubisco activation rate was identified on chromosome 7H. This is the first report on the identification of a QTL for rubisco activation rate *in planta* and the discovery opens the door to marker assisted breeding to improve whole canopy photosynthesis of barley. Further strength is given to this finding as this QTL colocalised with QTLs identified for steady state photosynthesis and stomatal conductance. Several other distinct QTLs were identified for these steady state traits, with a common overlapping QTL on chromosome 2H, and distinct QTLs for photosynthesis and stomatal conductance identified on chromosomes 4H and 5H respectively. Future work should aim to validate these QTLs under field conditions so that they can be used to aid plant breeding efforts.

**Highlight:** Significant variation exists in the photosynthetic induction response after a switch from moderate to saturating light across a barley doubled haploid population. A QTL for rubisco activation rate was identified on chromosome 7H, as well as overlapping QTLs for steady state photosynthesis and stomatal conductance.

## Introduction

By 2050, the global population is expected to rise to 9 billion and to meet future food demand we will need to increase crop production worldwide by 70% (Paul et al., 2019). Recent progress has been hindered by stagnating rates of annual yield increase, therefore novel breeding targets to improve crop yield potential are urgently needed. Improving the photosynthetic efficiency of crop species has now been shown to boost plant growth under field conditions (Kromdijk et al., 2016; South et al., 2019). Whilst these studies used genetic transformation to achieve such gains, they have proven that significant photosynthetic gains are possible in the field and that these can contribute to plant growth and crop yield. It is now imperative that we identify natural variation in photosynthetic traits in diverse populations and harness this variation through marker-assisted plant breeding techniques (Furbank et al., 2020).

Improving photosynthetic efficiency in dynamic environments has recently been highlighted as a key target to increase whole canopy carbon assimilation (Murchie et al., 2018). The light environment of the lower canopy is subject to continuous and dynamic change across the course of a day, caused by movement of the sun across the sky, sporadic cloud cover and/or movement of upper elements in the canopy caused by wind (Slattery et al., 2018). These processes can cause a leaf in low/moderate light one moment to suddenly be exposed to saturating light conditions the next. Often these short periods of direct sunlit illumination (referred to herein as ‘sunflecks’) only last a short period of time, in the order of seconds to minutes, yet they can account for as much as 90% of the daily accumulated light of lower canopy leaves (Pearcy, 1990). Photosynthesis under these dynamic light conditions is highly inefficient. Specifically, the rates of stomatal opening and activation of rubisco upon transition from low to high light significantly limit carbon assimilation. Interactive effects of several environmental factors on stomatal aperture are common in the field (Zeiger & Zhu, 1998; Talbott et al., 2003; Wang et al., 2008), meaning that stomata are operating in an integrated and hierarchical manner in response to multiple environmental stimuli (Lawson & Blatt, 2014). Stomatal responses to fluctuating light are therefore considered to be a much more challenging target for improvement than the biochemical limitations to photosynthesis (for a comprehensive review of limitations to dynamic photosynthesis see Kaiser et al., 2019).

Improving rubisco activation rate could be the low hanging fruit that allows plant breeders to boost whole canopy photosynthesis with few associated costs, specifically in terms of water and nutrient use. This is critical for a future where global environmental change is predicted to leave agricultural systems exposed to more frequent and more extreme drought and heat events. It has been estimated that we could increase daily carbon gain by as much as 21% in wheat if rubisco activation was instantaneous (Taylor & Long, 2017). Variation in rubisco activation kinetics has now been observed in crop species including soybean (Soleh et al., 2017), rice (Acevedo-Siaca et al., 2020) and wheat (Salter et al., 2019), and work with other species has indicated specific molecular targets and pathways that could accelerate rubisco activation speed, with a particular focus on rubisco’s catalytic chaperone rubisco activase (Rca) (in *Arabidopsis thaliana*, Mott et al., 1997; in *Nicotiana tabacum*, Hammond et al., 1998; and in *Oryza sativa*, Yamori et al., 2012). In our recent work with wheat, we found that by increasing the rate of rubisco activation of the slowest genotype included in our study to that of the fastest, daily carbon assimilation could be increased by 3.4% (Salter et al., 2019). However, in this work only ten genotypes of wheat were studied, the potential for improvement would likely be far more substantial if we were to investigate variation in this trait across a whole breeding population, and greater still if we were to investigate diversity within cultivars from diverse geographic locations or landraces.

Most recent studies of photosynthetic induction have tended to adopt the so-called ‘dynamic *A*/*c*i’ method (Taylor & Long, 2017; Salter et al., 2019), in which photosynthetic induction curves are measured at a number of different CO_2_ concentrations, allowing for the reconstruction of *A*/*c*_i_ curves throughout the induction response. Whilst this technique yields important fundamental data on the biochemical limitations during photosynthetic induction (most importantly rubisco carboxylation capacity, *V*_cmax_; and potential electron transport rate, *J*), it takes a very long time (> 6 hours per plant) and thus limits its use in large scale screenings for photosynthetic induction traits. Conversely, other techniques that yield less detailed information about the underlying physiology (such as that used by Soleh et al., 2016) take far less time (< 1 hour) and may be much more suitable for large-scale screenings of diverse plant material. The method of Soleh et al. (2016) involves measuring a single photosynthetic induction curve at a low intracellular CO_2_ concentration (i.e. < 300 μmol mol), at which it can be assumed that photosynthetic biochemistry is limited by rubisco rather than by electron transport. This allows for the reliable estimation of rubisco activation rate, with data comparable to those obtained using the ‘dynamic *A*/*c*_i_’ method (Taylor & Long, 2017).

We hypothesize that variation in photosynthetic induction kinetics may be inadvertently confounding efforts to improve steady state properties of photosynthesis. Steady state measurement techniques [such as spot measurements of photosynthesis (*A*) and stomatal conductance (*g*_s_), CO_2_ response curves and light response curves] all rely on the assumption that the leaf is fully acclimated to saturating light, and other environmental conditions inside the leaf chamber of the system, prior to measurement. Thus, these methods require a delay for the leaf to become equilibrated to the conditions inside the leaf chamber of the gas exchange system (referred to herein as the equilibration time). Although it is quite well established that adequate equilibration time is required for accuracy of steady state gas exchange measurements, an increasing demand for faster, higher throughput measurement techniques (Furbank & Tester, 2011) may make researchers complacent. Yet, few studies have quantitatively assessed the potential implications that may result from premature assumptions of steady state conditions, for instance, the identification of false quantitative trait loci (QTLs).

There is now compelling evidence that suggests whole canopy photosynthesis could be improved by harnessing natural variation in rubisco activation rate that exists across genotypes of crop species. However, no study to date has investigated or performed trait dissection for rubisco activation in a segregating mapping population. We sought to identify and characterise genetic variation in rubisco activation rate across a barley (*Hordeum vulgare* L.) doubled haploid (DH) mapping population *in planta* using gas exchange techniques. We then used chromosome interval mapping to identify QTLs and closely associated molecular markers. We were also interested in assessing whether false positive and/or false negative QTLs would be identified for “steady-state” photosynthetic properties (*A* and *g*_s_) if equilibration times were not long enough for steady state conditions to be reached.

## Methods

### Plant material and growth conditions

A DH barley (*H. vulgare* L.) population was obtained from a cross between the Australian barley cultivars Yerong and Franklin (Y/F). This population contained 177 DH lines and was maintained at the Plant Breeding Institute at The University of Sydney. The Y/F mapping population has been extensively used for QTL mapping for both morphological (Xue et al. 2008) and physiological (Zhang et al. 2016) traits, as well as disease resistance (Singh et al. 2014; Dracatos et al. 2016). In this study 131 lines from this population were phenotypically assessed for steady state and dynamic photosynthetic traits. Due to the availability of seed and genotypic data, only 127 DH lines were used for QTL analyses. A second DH barley population (from a cross between VB9104 and Dash) was also phenotyped for photosynthetic traits however due to the low number of lines with available genotypic data this population was not included in further analyses (phenotyping results are however presented in Figures S4 and S5).

Plants were grown in a controlled environment room for approximately five weeks prior to measurement. Day temperature was 25°C during a 14 h light period and night temperature was 17°C during a 10 h dark period. Relative humidity was maintained at 70% while daytime PPFD was approximately 600 μmol m^−2^s^−1^ at the top of the plants. Seeds were planted in potting mix enriched with slow-release fertilizer (Osmocote Exact, Scotts, Sydney, NSW, Australia). Six seeds per genotype were sown in 6 L pots and grown for three weeks before being thinned to three plants per pot. Seed was sown sequentially in time to make sure that all measurements were conducted at the same growth stage. Plants were watered daily to field capacity.

### Photosynthetic measurements

Plants were moved from the controlled environment room to a temperature-controlled growth cabinet [temperature 25°C; relative humidity 70%]. Two or three of the youngest fully expanded leaves of a single plant were sealed in a 2×6 cm leaf cuvette (Li6400 11; LI-COR, Lincoln, NE, USA) fitted to a LI-COR LI-6400XT gas exchange system to fill the cuvette without overlapping. This simulated an instantaneous shift in light intensity from 600 μmol m^−2^ s^−1^ to 1300 μmol m^−2^ s^−1^, similar to the conditions experienced by a lower canopy leaf during a sunfleck. Chamber conditions were set to closely match those of the controlled environment room [leaf temperature 25°C; cuvette CO_2_ (*C*_a_) 400 µmol mol^−1^; relative humidity 70%], with the exception of PPFD which was set to 1300 µmol m^−2^ s^−1^ using a red-green-blue light source (Li6400 18A; LI-COR) set to 10% blue and 90% red light. Measurements of photosynthetic gas exchange rates (*A* and *g*_s_) were recorded once per minute immediately after the leaf was inserted into the chamber until photosynthesis had reached steady state. Preliminary photosynthetic light response curves were measured with plants grown under the same conditions to ensure that 1300 μmol m^−2^ s^−1^ was saturating and that 600 μmol m^−2^ s^−1^ was non-saturating (results shown in Figure S1).

Rubisco activation rate was calculated using a modified method of Soleh et al. (2016). Photosynthetic data was first normalised to an assumed intercellular CO_2_ concentration (*c*_i_) of 300 µmol mol^−1^, using the following equation:

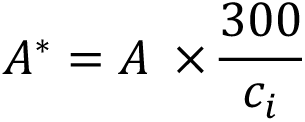

where *A*^*^ is the normalised photosynthetic rate, *A* is the measured photosynthetic rate and *c*_i_ is the measured intercellular CO_2_ concentration. This effectively removed the influence of stomatal opening/closure for the induction phase. The initial rubisco activation rate (1/τ) was modelled from the plot of the logarithmic difference between *A*^*^ and its maximum value after induction (*A*^*^_max_) against the time taken for induction (representative data shown in Figure 1). From this plot, the value of 1/τ was determined from the slope of the linear regression on data points in the range of 2 to 5 mins after induction, and points after this that aligned well with these initial points (with an R^2^ > 90%).

### Genetic analysis and QTL mapping

The genotypic data and genetic linkage map for the Yerong/Franklin DH population used for QTL analysis for rubisco activity and steady state photosynthetic traits in the present study was previously described by Singh et al. (2015). In brief, the Y/F genetic map is comprised of 496 DarT and 28 microsatellite (SSR) markers spanning 1,127cM across all seven chromosomes, 1H to 7H (Wenzl et al., 2006).

**Figure 1.**
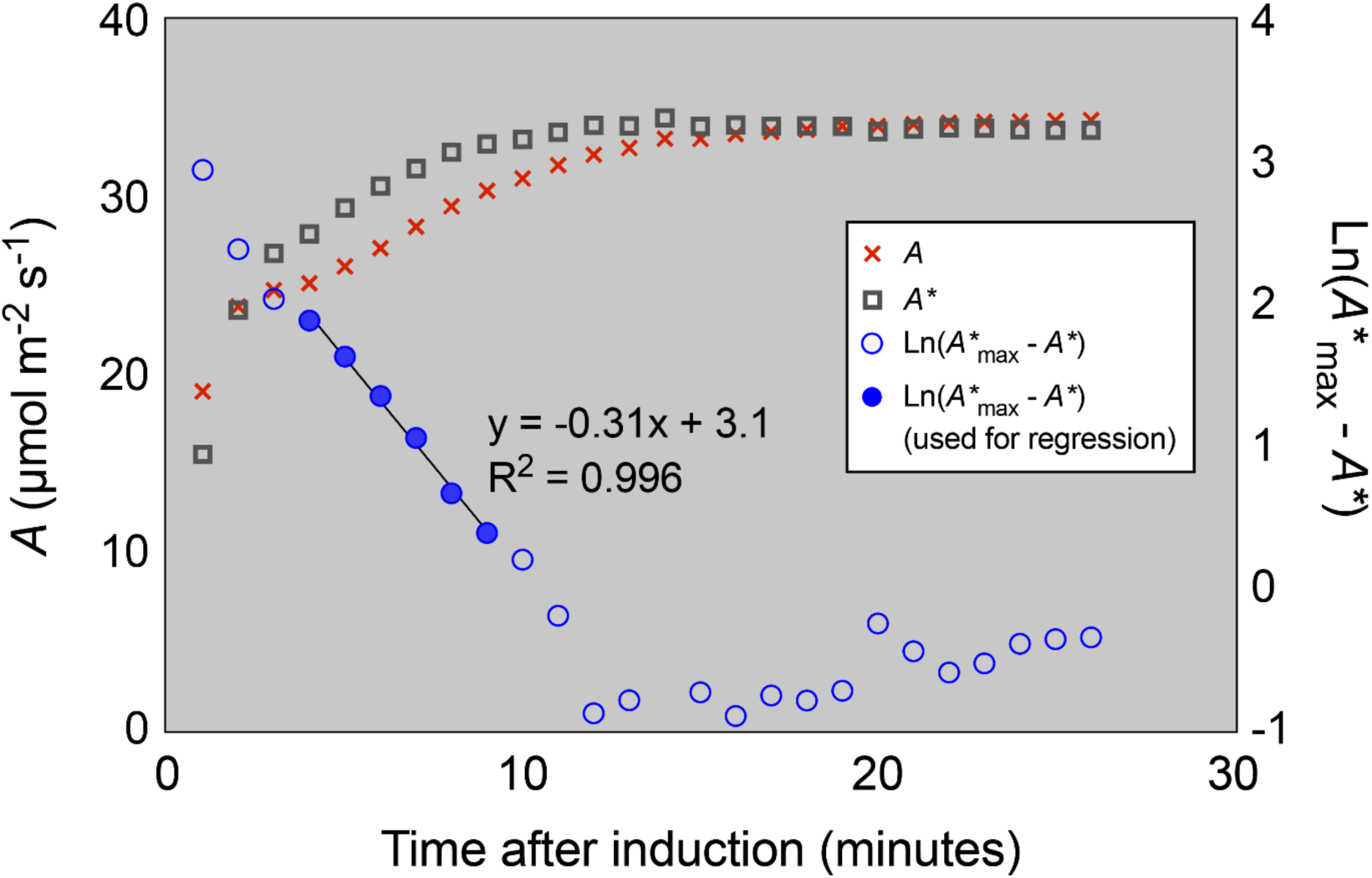
Example of a typical leaf photosynthetic induction response and the linear regression used to calculate rubisco activation rate (1/τ). The orange crosses represent the measured photosynthetic rate *A*; the grey squares the *c*_i_ = 300 µmol mol^−1^ normalised photosynthetic rate *A*^*^; and the blue circles the logarithmic difference between the fully induced photosynthetic rate *A*^*^_max_ and *A*^*^. Filled circles represent the data points used in the linear regression to estimate 1/τ, the rubisco activation rate. The slope of the regression represents 1/τ, in this case 0.31 min^−1^.

A subset of 127 lines were selected for QTL mapping analysis. Markers were selected every 10 cM so that the whole genome was evenly covered. Composite interval mapping (CIM) methods were used in QTL Cartographer version 2.5 (North Carolina State University, Raleigh, NC, USA), carrying out 1,000 iterations permutation analysis with steps at 1 cM, and with a 0.05 confidence level for all traits.

### Statistical analyses

All other modelling and statistical analyses were performed in R (R Core Team, 2019).

## Results

### Photosynthetic induction kinetics

While specific photosynthetic induction kinetics were found to vary across individual leaves and genotypes, general trends were quite clear (representative induction curves from one day of measurements is shown in Figure 2). Net photosynthesis (*A*) increased immediately after transition from to saturating light for all leaves. Stomatal responses were more variable than those of photosynthesis but there tended to be an initial reduction in *g*_s_ after transition to saturating light and then a gradual rise towards steady state. By normalising photosynthesis to a constant *c*_i_ of 300 ppm, we were able to obtain a measure of photosynthesis limited by rubisco carboxylation unobstructed by variation in stomatal kinetics (*A*\*). *A*\* showed a similar trend to *A*, increasing immediately after the switch from low to high light.

**Figure 2.**
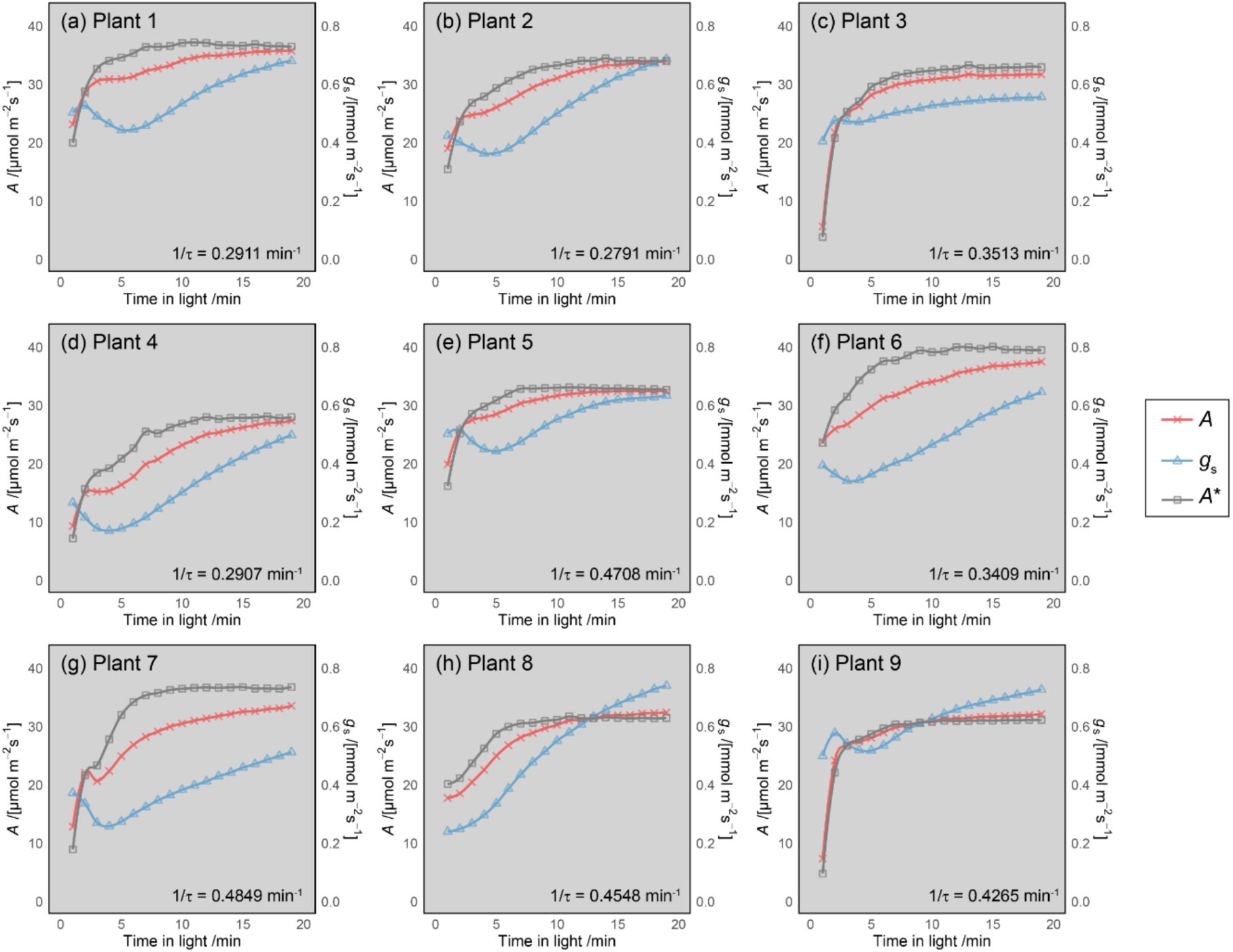
Induction curves for net photosynthesis, *A* (red crosses); stomatal conductance, *g*_s_ (blue triangles); and *c*_i_ = 300 ppm normalised photosynthesis, *A\** (grey squares), after a switch from moderate to saturating light. Data shown in panels (a) – (i) are representative induction curves from one day of measurements in individual plants. The value of 1/τ is shown in each panel for reference.

### QTLs for rubisco activation rate

Rubisco activation rates of the parental lines Yerong and Franklin were found to differ, with within-genotype medians of 0.38 min^−1^ and 0.74 min^−1^ respectively. Wide variation in 1/τ was found across the population (Figure 3), with within-genotype medians ranging from 0.099 min^−1^ to 0.74 min^−1^. Interestingly, the parental line Franklin was found to have the fastest rate of rubisco activation. A frequency distribution of 1/τ was plotted for the population and was found to follow a normal distribution suggesting that rubisco activation rate was under complex genetic control (Figure S2). CIM analysis revealed the presence of a distinct QTL for rubisco activation rate (Figure 4; further details in Table 1). Q1/τ.sun-7H, was located at 41.67 cM on chromosome 7H (proximal to DarT marker bPb-9601 marker) accounting for 10.48% of the phenotypic variance in this trait.

**Table 1.** QTLs for dynamic and steady state photosynthetic traits identified in the mapping population.

| Trait | QTL | Chromosome | Position (cM) | Nearest Marker | Explained variance (%) | Additivity | LOD |
| --- | --- | --- | --- | --- | --- | --- | --- |
| **1/τ** | Q1/τ.sun-7H | 7H | 41.67 | bpb-9601 | 10.48 | -0.07 | 4.40 |
| **A** | QA.sun-2H | 2H | 27.03 | bpb-0003 | 9.20 | -3.85 | 4.31 |
| **A** | QA.sun-4H | 4H | 66.68 | bpb-2305 | 5.84 | -4.37 | 2.64 |
| **A** | QA.sun-7H | 7H | 41.67 | bpb-9601 | 10.80 | -4.30 | 5.18 |
| **g<sub>s</sub>** | Qg<sub>s</sub>.sun-2H | 2H | 35.91 | bpb-8750 | 11.80 | -0.09 | 5.41 |
| **g<sub>s</sub>** | Qg<sub>s</sub>.sun-5H | 5H | 53.39 | bpb-5532 | 6.49 | 0.07 | 2.98 |
| **g<sub>s</sub>** | Qg<sub>s</sub>.sun-7H | 7H | 41.28 | bpb-4989 | 13.75 | -0.11 | 6.80 |

**Figure 3.**
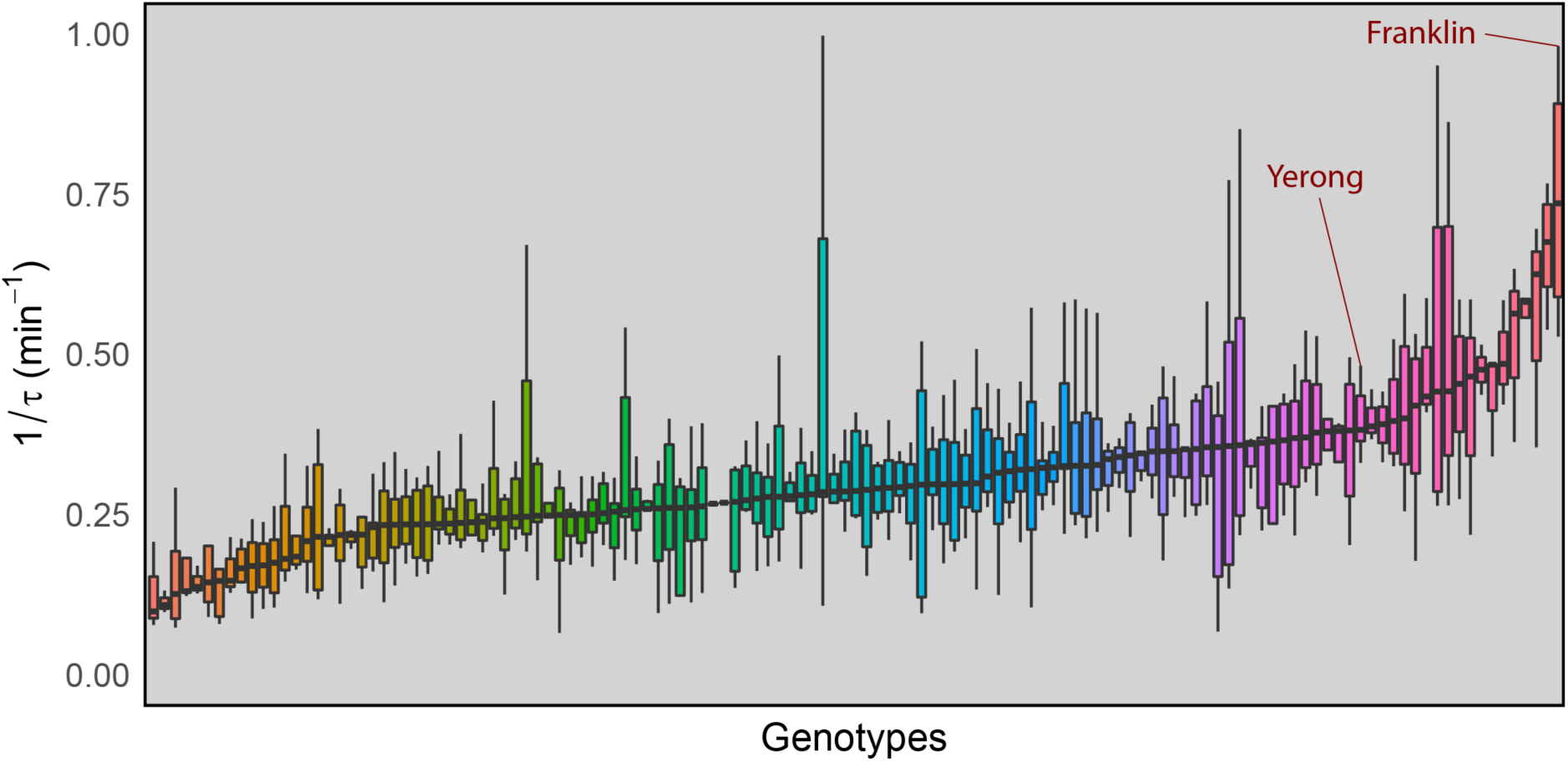
Distribution of rubisco activation rate (1/τ) across genotypes of the Yerong/Franklin DH population. Each bar represents a single genotype. Parental lines are highlighted. Colours are arbitrary.

**Figure 4.**
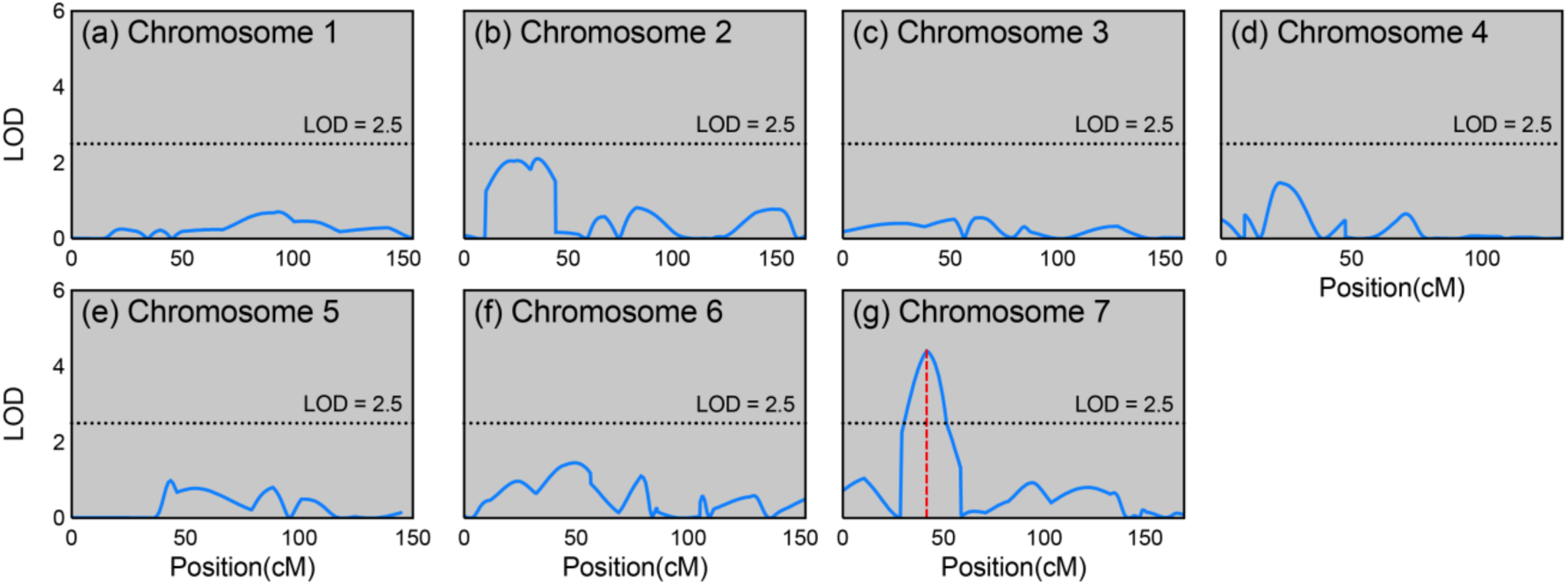
Logarithm of odds (LOD) traces from composite interval QTL mapping analysis for 1/τ. LOD values are plotted against the position on the chromosomes. The significance threshold LOD of 2.5 is indicated by the dotted line in each plot. Vertical dashed red lines represent identified QTLs.

### Steady state photosynthesis and equilibration time tests

Variation was also found in steady state photosynthetic rates across the population (Figure 5). Median rates of *A* and *g*_s_ were 17.45 μmol m^−2^ s^−1^ and 0.31 mmol m^−2^ s^−1^, respectively. From this phenotyping data, there was no correlation found between steady state *A* and 1/τ (p > 0.05; Figure 6).

**Figure 5.**
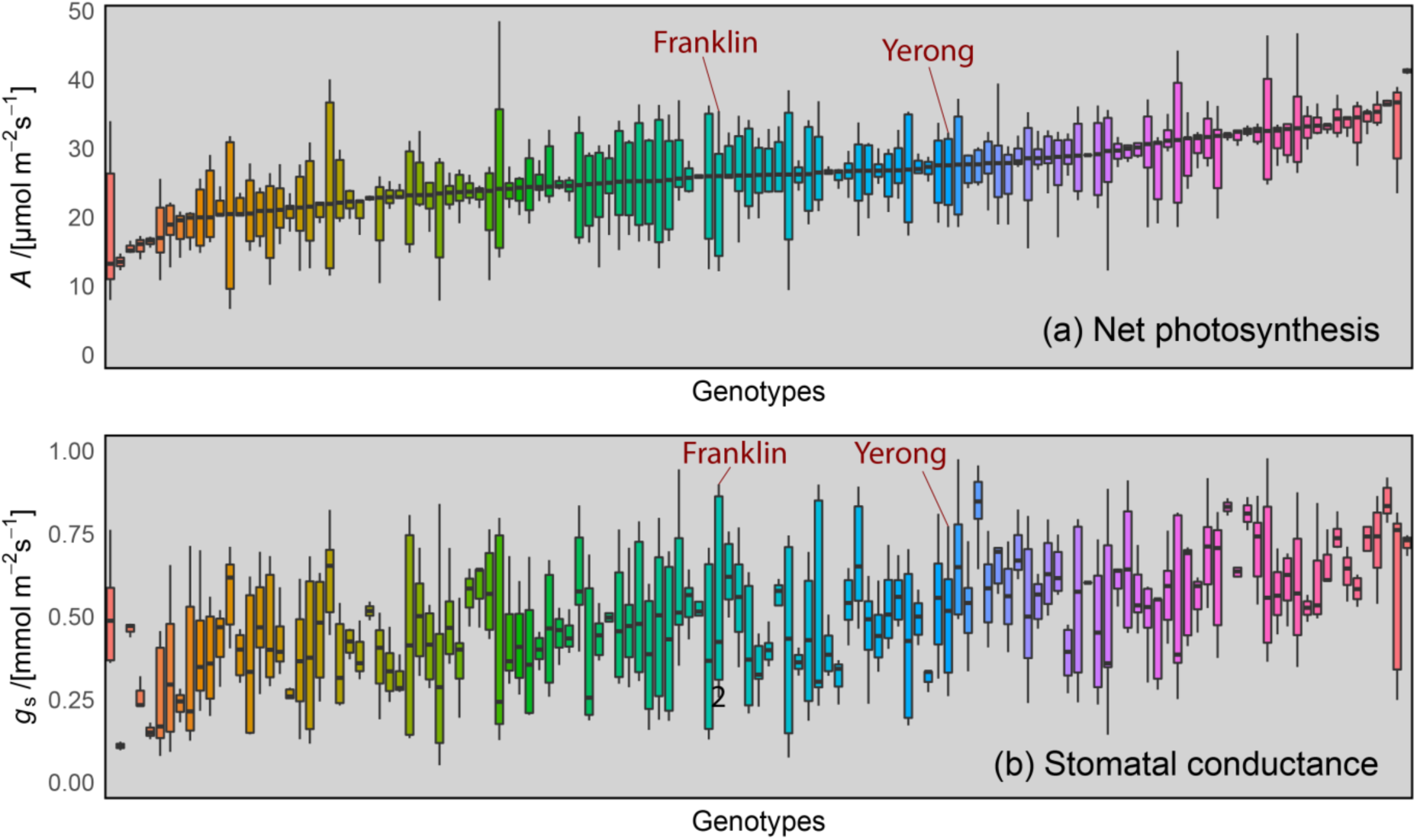
Distribution of steady state (a) *A* and (b) *g*_s_ across genotypes of the Yerong/Franklin population. Each bar represents a single genotype. Parental lines are highlighted. Note that colours are arbitrary but are consistent for genotypes in panels (a) and (b).

**Figure 6.**
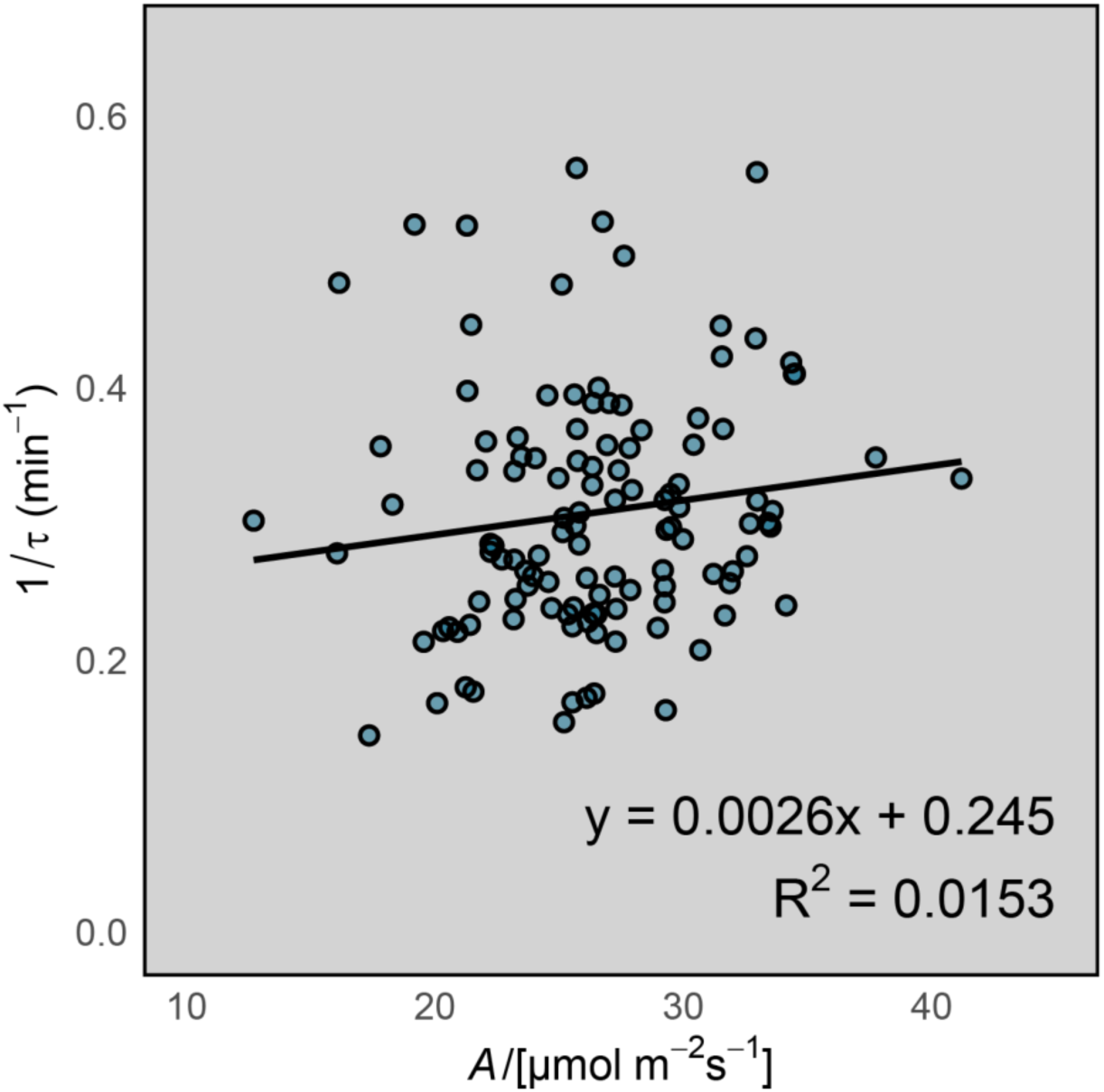
Relationship between steady state *A* and 1/τ. Each point represents a genotype. Values are genotype means.

As hypothesized, “steady-state” photosynthetic rates were substantially underestimated if measurements were recorded without sufficient equilibration time (Table 2). This was more pronounced the earlier the measurements were recorded after enclosing the leaf in the chamber of the IRGA. Mean values of *A* and *g*_s_ were both underestimated by 21% at five minutes compared to steady state. It should be noted that although some of the fastest genotypes reached steady state after five minutes, most of the lines did not. In fact, *g*_s_ was underestimated by 82% for one of the genotypes and *A* was underestimated by 54% for another if measurements were recorded after just five minutes.

**Table 2.** Distribution features for photosynthetic rate (*A*) and stomatal conductance (*g*_s_) across the population at 5, 10 and 15 minutes into photosynthetic induction.

| Trait | Time point (minutes) | Minimum | 25% Percentile | Median | 75% Percentile | Maximum | Mean |
| --- | --- | --- | --- | --- | --- | --- | --- |
| $A$<br>($\mu\text{mol m}^{-2} \text{s}^{-1}$) | 5 | 7.1 | 16.5 | 20.0 | 24.9 | 36.5 | 20.4 |
|  | 10 | 9.6 | 20.0 | 23.3 | 26.9 | 40.2 | 23.4 |
|  | 15 | 12.2 | 21.5 | 24.9 | 28.4 | 41.8 | 25.0 |
| $g_s$<br>($\text{mmol m}^{-2} \text{s}^{-1}$) | 5 | 0.044 | 0.259 | 0.365 | 0.522 | 0.947 | 0.397 |
|  | 10 | 0.069 | 0.294 | 0.392 | 0.508 | 0.947 | 0.412 |
|  | 15 | 0.107 | 0.330 | 0.441 | 0.544 | 0.940 | 0.451 |

To assess the importance of equilibration time for accurate identification of steady state QTLs, QTL mapping was first performed for steady state *A* and *g*_s_. Frequency distributions were plotted for both traits and they followed a normal distribution suggesting they are under complex genetic control (Figure S3). Several QTL were found for both *A* and *g*_s_ (Table 1). Trait co-location was observed on chromosome 7H whereby the position of the Q1/τ.sun-7H QTL was almost identical to QTL for both *A* and *g*_s_. This suggests a region on the short arm of chromosome 7H either carries a single gene or more likely a cluster of genes responsible for the genetic control of photosynthesis, stomatal conductance and rubisco activation. For steady state *A* and *g*_s_, additional overlapping and distinct QTL were identified. A common overlapping QTL for both *A* and *g*_s_ was identified, peaking at 27.03 cM on chromosome 2H, whilst distinct QTL were identified on chromosomes 4H (41.67 cM) and 5H (53.39 cM) for *A* and *g*_s_ respectively.

QTL mapping was then performed with data collected at five, ten and fifteen minutes after the start of induction for comparison with detected steady state QTLs (coloured traces in Figure 7). Although most QTL were still identified with non-steady state data, the significance these QTL peaks were found to be weakened under non-steady state conditions. This was particularly evident for the *g*_s_ QTL identified on chromosome 7H (Figure 7h), with the LOD score of this QTL dropping from 6.8 when using steady state data to 4.9, 3.9 and 3.3 when using data collected at 15 min, 10 min and 5 min after the start of induction, respectively.

**Figure 7.**
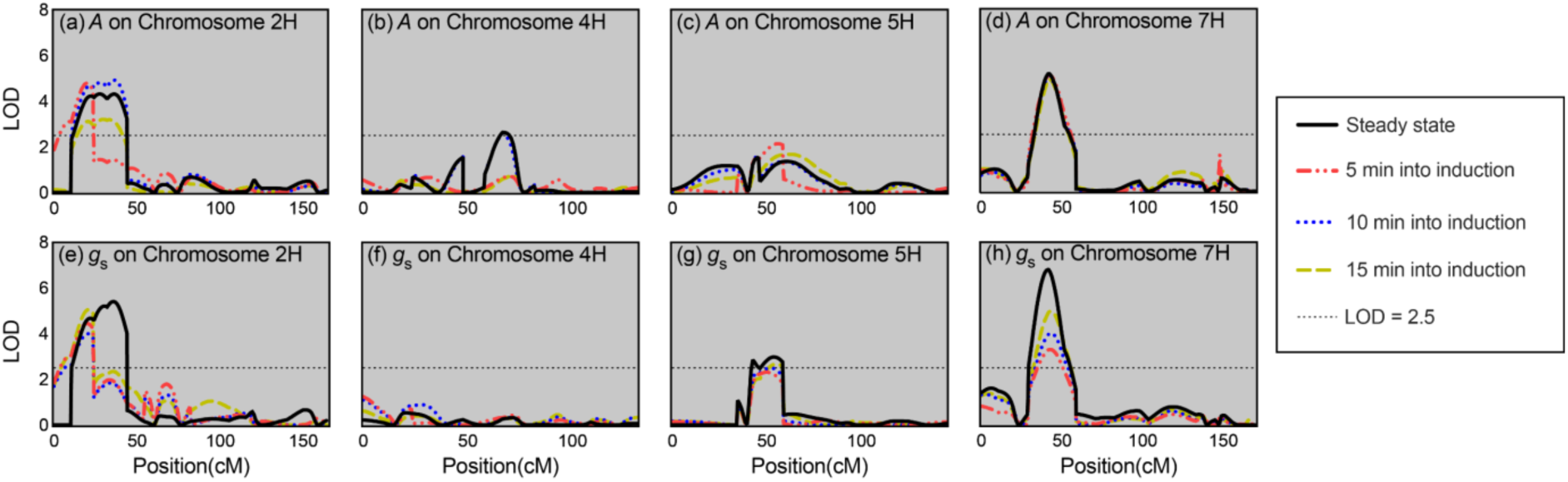
Logarithm of odds (LOD) traces from composite interval QTL mapping analysis for *A* and *g*_s_ in the Yerong/Franklin DH population. LOD values are plotted against the cM position on chromosomes 2H, 4H, 5H and 7H. The threshold LOD of 2.5 is indicated by the horizontal dotted line in each plot. Note that LOD plots for chromosomes 1H, 3H and 6H are not shown as there were no significant QTLs identified on these chromosomes.

## Discussion

We have identified QTLs for *in planta* rubisco activation rate for the first time in any species. As in other crops, we found rubisco activation rate to be a highly variable trait across genotypes of barley, aiding in the discovery of a significant QTL in our doubled haploid population. QTLs were also identified for steady state photosynthetic parameters, including co-localised QTLs for *A*, *g*_s_ and 1/τ on chromosome 7H. The importance of adequate equilibration time in the measurement of steady state gas exchange was highlighted by comparing these results to those obtained using arbitrary non-steady state rates at 5, 10 and 15 min after the start of induction. The significance of QTLs was reduced if steady state conditions had not been reached.

### Improving whole canopy photosynthesis

It is well established that improving photosynthesis has the potential to increase crop yield (for a review of recent progress see Simkin et al., 2019). However, until now research has invariably focussed on only the uppermost leaves of the canopy under optimal conditions (i.e. continuous saturating light, 25°C). This approach has its merits because these leaves have the most light available to them and their contribution to whole canopy photosynthesis reflects this (Osborne et al. 1998). Yet, for monocot cereal species such as wheat and barley there have been few studies that have shown flag leaf photosynthesis to correlate well with crop yield (Richards et al., 2000). Whole canopy photosynthesis, and more specifically the cumulative rate of photosynthesis over the growing season, can be a much more reliable determinant of crop yield (Wu et al., 2019). Accordingly, there has been a recent shift in research focus towards dynamic photosynthetic traits. This is important because whilst some studies have found weak relationships between steady state and dynamic photosynthetic traits (Salter et al., 2019) other studies have not found any relationship (Soleh et al, 2017; Acevedo-Siaca et al., 2020). Our results also showed no link between steady state *A* and 1/τ (Figure 6), although the co-localisation of QTLs on chromosome 7H suggests they both may be controlled by the action of a single gene or a cluster of closely linked genes at the same chromosomal location.

Significant improvements in photosynthesis and resultant increases in plant growth have now been achieved under field conditions through genetic modification of model plant species (15% increased biomass production by accelerating recovery from photoprotection, Kromdijk et al., 2016; and 40% increased biomass via engineering of a photorespiratory bypass, South et al., 2019) and recent modelling has highlighted the potential of improving several dynamic photosynthetic traits on whole canopy photosynthesis (Wang et al., 2020). It is important now that we explore and exploit natural variation in photosynthetic traits across plant populations (for review see Furbank et al., 2020). As in previous studies with other species, we identified significant variation in rubisco activation rate across barley genotypes. We identified a QTL for rubisco activation rate, as well as several QTLs for steady state *A* and *g*_s_. Q1/τ.sun-7H was flanked by the bpb-9601 DArT marker which has previously been associated with both grain yield and crop spike number in the Yerong/Franklin population (Xue et al., 2010). This marker is of particular interest as it also flanks QTLs that we identified for steady state *A* and *g*_s_ (Q*A*.sun-7H and Q*g*_s_.sun-7H in Table 1), highlighting its utility for marker assisted selection (MAS). MAS exploiting natural variation between barley genotypes can now be achieved through the development of a high throughput codominant marker using the sequence information from the closely associated bpb-9601 DArT marker identified in this study. MAS for both steady-state and dynamic photosynthetic traits in barley now provides potential to improve daily photosynthetic carbon gain in both sporadically sunlit lower canopy and fully sunlit upper canopy leaves, bolstering whole canopy photosynthesis and contributing to yield potential.

Whilst we observed segregation for three different photosynthetic traits in the barley mapping population studied, and the V/D population presented in Figures S4 and S5, these populations were not specifically developed to investigate photosynthesis. Future work in this area would hugely benefit from phenotyping a diverse panel of barley accessions to either develop additional trait-specific mapping populations using parents with contrasting photosynthetic properties or use a genome wide association scan (GWAS) approach to mine for novel favourable alleles based on natural variation in photosynthetic traits. This may also include the investigation of crop wild relatives (Castañeda-Álvarez, 2016). Such approaches have already yielded promising outcomes for other desirable traits in crop species, including salinity (in barley, Saade et al., 2016) and drought tolerance (Venuprasad et al., 2009).

Due to the recent availability of multiple reference genomes for cultivated and wild barley, the precision of GWAS studies and ability to rapidly clone genes of interest from cereal crops is continually improving. Further studies are required to determine whether each of the traits studied are under control by a single gene or more complex genetic control within the QTL region on chromosome 7H. Further mendelisation of the 7H QTL by intercrossing select DH lines from the Y/F population will enable the development of a large <u>segregating</u> F_2_ fine-mapping population for positional cloning of the 7H QTL to unravel the underlying genetic and biological mechanisms involved.

Our study focussed on a step change from moderate (600 µmol m^−2^ s^−1^) to saturating light (1300 µmol m^−2^ s^−1^), rather than low to high light as has been reported previously (i.e. 50 – 1500 µmol m^−2^ s^−1^ in Taylor and Long, 2017). We felt this approach would provide more valuable information for plant breeding, as it more accurately represents the light regime experienced by the second youngest leaves in the canopy, which for wheat have been reported to receive between 300 – 700 µmol m^−2^ s^−1^ PPFD when not in a sunfleck (Townsend et al., 2018). Whilst leaves lower in the canopy receive much less light than this (< 300 µmol m^−2^ s^−1^), these leaves are also less likely to be exposed to sunflecks and also have a much-reduced photosynthetic capacity (Townsend et al. 2018), so contribute considerably less to whole canopy photosynthesis. Our results show that rubisco activation rates after a switch from moderate to high light in barley (median 1/τ = 0.28 min^−1^) are similar to those that have been reported from low to high light in other species (0.3 – 0.45 min^−1^ in rice, Yamori et al., 2012; 0.24 – 0.42 min^−1^ in soybean, Soleh et al., 2016; and 0.25 – 0.33 min^−1^ in wheat, Taylor and Long, 2017), albeit with greater variation. It would therefore seem that the same biochemical processes, likely related to the amount of and form of rubisco activase present in the leaves (Carmo-Silva and Salvucci, 2013), are involved in photosynthetic induction under the two induction scenarios.

### Limitations and future directions

Our study has focussed on rubisco activation however this is only one part of the dynamic photosynthesis puzzle, in which all the pieces must be investigated to fully understand potential improvements that could be made to whole canopy photosynthesis. Responses of stomata can also limit photosynthesis in fluctuating light. Faster stomatal opening has now been shown to improve net photosynthesis and biomass production in overexpressing mutants of *Arabidopsis thaliana* compared to wild type plants (Kimura et al., 2020). And so, if improvements are made to rubisco activation rate without also considering rates of stomatal opening/closure, the dominant limitation will likely shift in the direction of the stomata. In effect, this could nullify any improvements made to rubisco activation in terms of net photosynthesis. On a positive note, recent work has highlighted that stomatal traits can be linked to rubisco kinetics during leaf development in some plant species (Conesa et al., 2019), and it has long been realised that stomata respond to photosynthetic activity in the mesophyll (Messinger et al., 2006). It is therefore conceivable that improving rubisco activation rate through targeted plant breeding could also inherently result in improved stomatal responses. Regardless, there is a definite need for future work in this area to address dynamic responses of stomata, rubisco and other biochemical processes (i.e. non-photochemical quenching) of photosynthesis together, rather than focussing on each in isolation.

In this study, we measured photosynthetic induction and identified associated QTLs in plants grown under optimal and controlled conditions. The next important step is for photosynthetic induction traits to be investigated in field grown plants with established canopies. Traditional gas exchange techniques combined with new higher throughput techniques based on thermography (for dynamic stomatal traits; Vialet-Chabrand & Lawson, 2020), hyperspectral imaging and chlorophyll fluorescence (for dynamic photosynthetic parameters; McAusland et al., 2019; Meacham-Hensold et al., 2020) may offer the potential to screen these two populations in the field and validate the QTLs we identified in this study. It is also important that we understand if these QTLS are strong under sub-optimal conditions (i.e. under drought or heat stress), as for most growers such conditions can be common during a growing season.

### A note on gas exchange methodology

It is common practice to allow a leaf to stabilise to the chamber conditions of an IRGA, yet the recent push for “high throughput” and “big data” approaches in plant physiology may have made researchers complacent. We hypothesized that this complacency could impact detected QTLs for photosynthesis and stomatal conductance, and indeed we found that using non-steady state rates (i.e. before leaves had equilibrated to chamber conditions) resulted in less accurate detection of QTLs. It is likely that false QTL identifications are worsened by the high variability in photosynthetic induction kinetics that exists across this population (and has also been found in other crop species) and the fact that there is no clear relationship between steady state and dynamic photosynthesis. This result reinforces the importance of good gas exchange technique. The push for high-throughput measurements has resulted in new fast methods, such as the Rapid *A*/*c*_i_ method (Stinziano et al., 2017), being developed yet it must be highlighted that most of these methods still rely on the assumption of steady state conditions and these will therefore still be limited by equilibration time.

We suggest that plant physiologists treat this as a methodological opportunity instead of a hindrance. Rather than just waiting for the leaf to reach steady state and then recording a point measurement or photosynthetic response curve, the photosynthetic induction phase could always be logged continuously as soon as the leaf enters the chamber. Not only would this provide extra data on photosynthetic induction, it would also provide transparency and confidence in the data. Specifically, the researcher and their peers would be able to backcheck to ensure that steady state conditions had been reached. In the past, technical limitations may have prevented such an approach, but new gas exchange instruments have both the computational power and environmental control to establish this as common practice.

## Conclusions

In this study, we found wide variation in photosynthetic induction to fluctuating light across a barley mapping population. This variation allowed us to identify a QTL for rubisco activation rate, the position of which overlapped QTLs for steady state photosynthesis and stomatal conductance. These QTLs lie close to molecular markers that could be used for selection in plant breeding programs. Future work should aim to validate these QTLs under field conditions so that they can be used to aid plant breeding efforts.

## Acknowledgements

This work was funded by the Grains Research and Development Corporation (contract US00056). WTS was supported by the Australian Research Council (ARC), Industrial Transformation Research Hub —Legumes for Sustainable Agriculture (IH140100013) and the Grains Research and Development Corporation. The authors thank Dr Meixue Zhou, University of Tasmania for the available genotypic data and Dr Davinder Singh and Professor Robert Park for maintenance of seed of the doubled haploid barley populations.

## Supplementary figures

**Figure S1.**
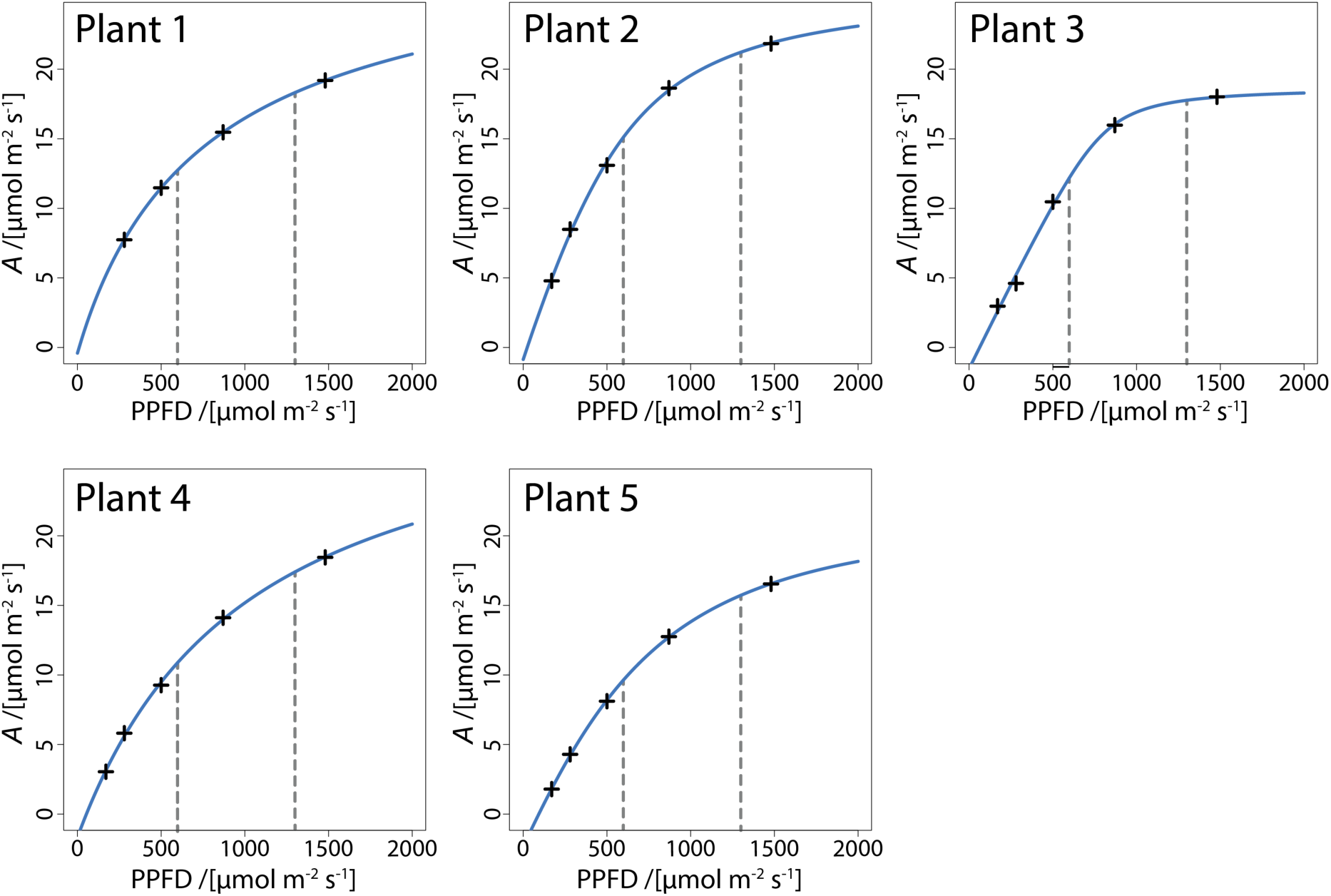
Photosynthetic light response curves measured on plants of the parental line Dash grown under the same growth conditions as experimental plants. Curves were fitted to a non-rectangular hyperbola model using non-linear least squares in R (*nls*; R Language and Environment) as per Salter *et al*. (2019). Vertical dashed lines are shown at 600 μmol m^−2^ s^−1^ and 1300 μmol m^−2^ s^−1^ to highlight the moderate to high light induction phase measured in this study.

**Figure S2.**
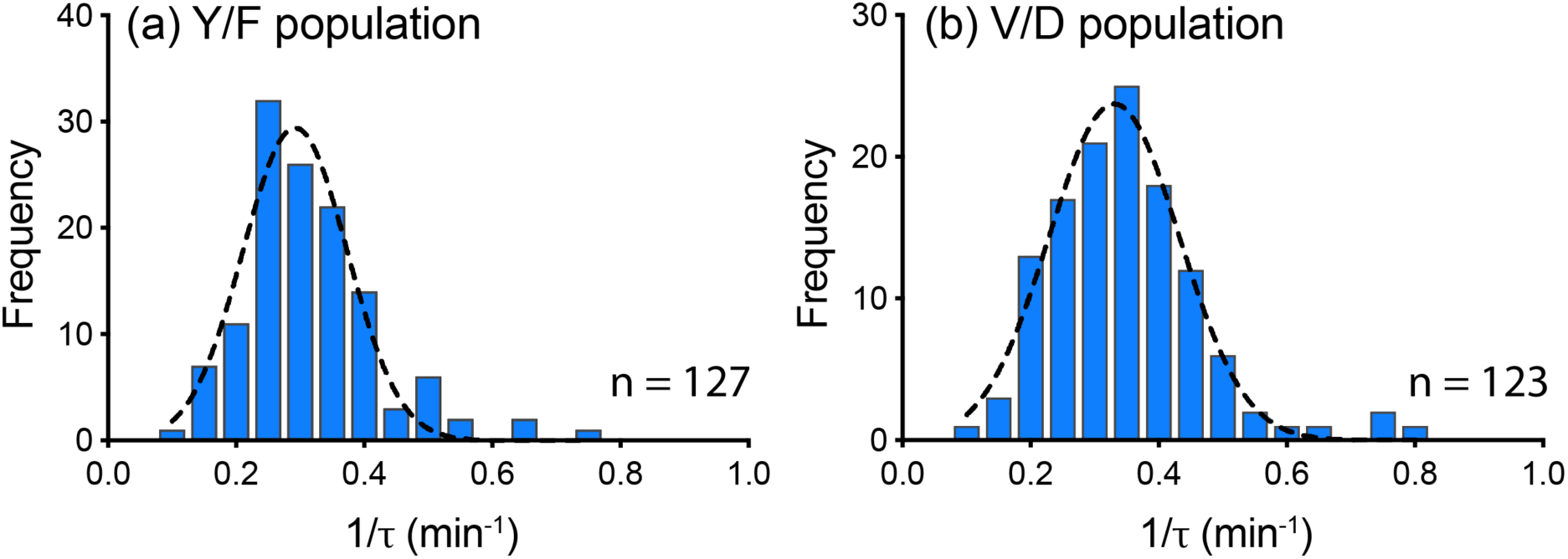
Frequency distributions of 1/τ for the Y/F DH population.

**Figure S3.**
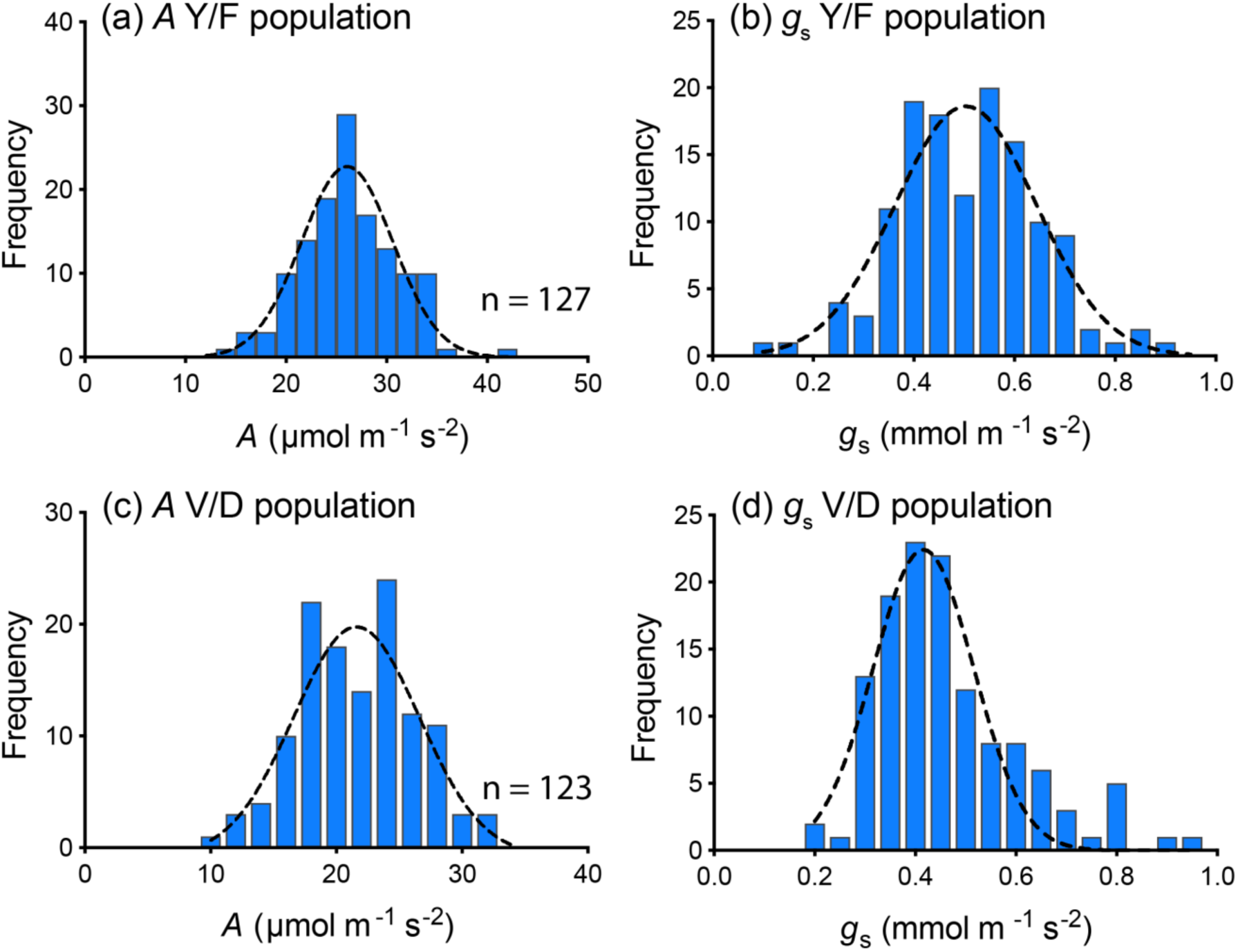
Frequency distributions of steady-state (a) *A* and (b) *g*_s_ for the Y/F DH population.

**Figure S4.**
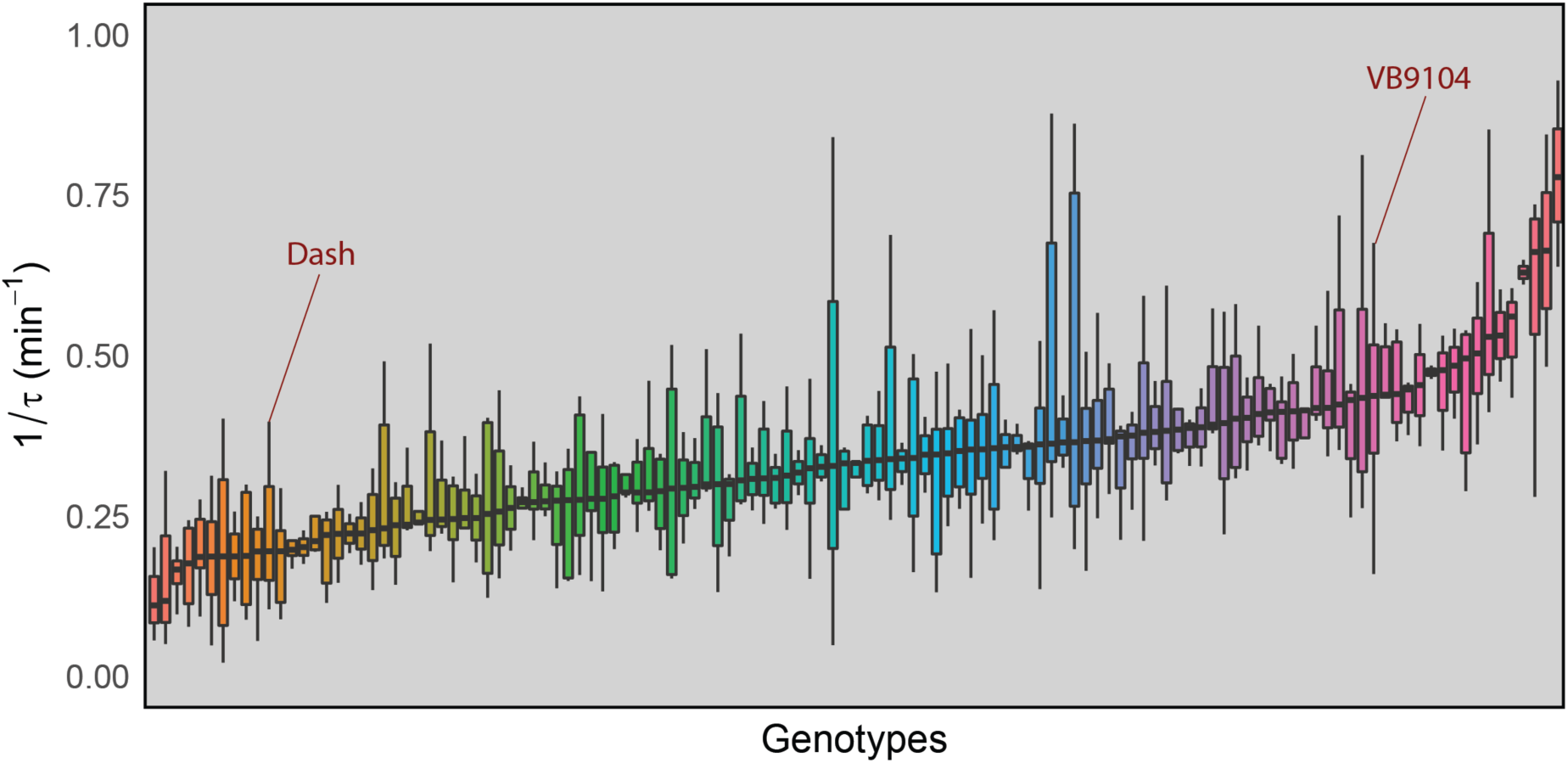
Distribution of rubisco activation rate (1/τ) across genotypes of the V/D DH population. Each bar represents a single genotype. Parental lines are highlighted. Colours are arbitrary.

**Figure S5.**
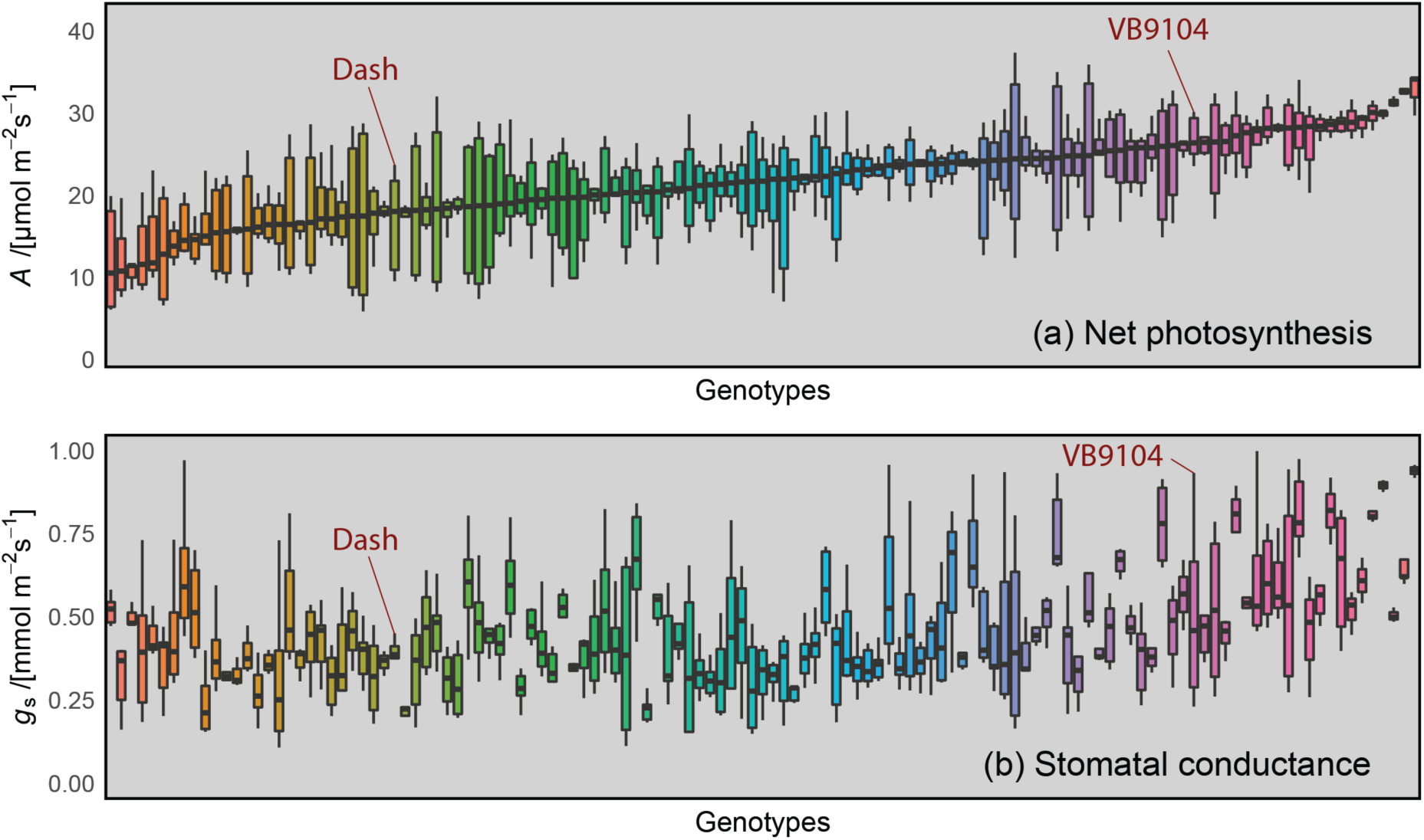
Distribution of steady state (a) *A* and (b) *g*_s_ across genotypes of the VB9104/Dash population. Each bar represents a single genotype. Parental lines are highlighted. Note that colours are arbitrary but are consistent for genotypes in panels (a) and (b).

## List of supplementary files

*FileS1.xlsx –* Yerong/Franklin dynamic and steady state gas exchange phenotypic data.

*FileS2.xlsx –* Results of composite interval mapping of dynamic and steady state photosynthetic traits.

